# Activation of neuromodulatory axon projections in primary visual cortex during periods of locomotion and pupil dilation

**DOI:** 10.1101/502013

**Authors:** Rylan S. Larsen, Emily Turschak, Tanya Daigle, Hongkui Zeng, Jun Zhuang, Jack Waters

## Abstract

Neuromodulators such as acetylcholine, noradrenaline (norepinephrine), and serotonin are released into the cortex by axons ascending from subcortical nuclei. These neuromodulators have been hypothesized to influence cortical function during behavioral periods such as arousal, locomotion, exploration, and attention. To determine when these neuromodulatory projections were active, we expressed the genetically-encoded calcium sensor GCaMP6 in neuromodulatory axons which project to the mouse primary visual cortex and performed two-photon microscopy to monitor their activity *in vivo*. We observed that the fluorescence of both cholinergic and noradrenergic axons increased during periods of pupil dilation, with the fluorescence of the axons rising less than one second before eye pupil dilation. We also observed increases in cholinergic and noradrenergic axon fluorescence periods of locomotion, which was accompanied by pupil dilation and nasal (forward) movement of both pupils. Locomotion was preceded by a rise in axonal fluorescence with a timing and amplitude that matched the subsequent pupil dilation, but axon fluorescence was more sustained than expected from the pupil dilation, suggesting that there is an additional physiological factor that affects cholinergic and noradrenergic axon activity in primary visual cortex during locomotion.

## INTRODUCTION

Changes in neuronal network activity correlate with periods of arousal, locomotion, exploration, and attention (Lee and Dan, 2012). These brain states are also accompanied by physiological changes such as locomotion, pupil dilation and eye movements (Larsen and Waters, 2018). Changes in brain state also influence the cortical processing of sensory stimuli. For example, the spiking of neurons in visual cortex in response to visual stimuli is more reliable and selective during periods of pupil dilation (Reimer *et al*., 2014), periods of locomotion are associated with an increase in stimulus discriminability (Dadarlat and Stryker, 2017), and the signal to noise ratio from individual neurons is higher following an arousing stimulus (Vinck *et al*., 2015). These effects on visual processing are thought to be mediated, at least in part, by changes in the signaling from subcortical neuromodulatory inputs that are activated during changes in brain state (McGinley et al., 2015b).

The mouse neocortex receives axonal projections from cholinergic, noradrenergic, serotonergic and, in anterior cortex, dopaminergic neuromodulatory neurons (Bjorklund and Dunnett, 2007; Ren et al., 2018; Schwarz and Luo, 2015). These projections have long been hypothesized to drive the orchestrated changes in cortical network activity that accompany a shift in behavioral state (Fu et al., 2014; Kalmbach et al., 2012; Kalmbach and Waters, 2014; McGinley et al., 2015a; Roerig et al., 1997). Until recently, observing behaviors in which each of these neuromodulatory systems influences a local region of cortex was difficult. Previous studies instead attempted to infer the effect of a neuromodulation on the cortex from somatic recordings or broad optogenetic or chemogenetic manipulation of a population of neuromodulatory neurons without regard to potential heterogeneity in their cortical projection target (Carter et al., 2010; Harrison et al., 2016; Pinto et al., 2013). Recent advances in viral tracing have demonstrated that neuromodulatory neurons of a given neurotransmitter type can both receive diverse inputs and have distinct projection targets, despite having the same neurotransmitter-defined identity (Kim et al., 2016; Obermayer et al., 2017; Ren et al., 2018; Schwarz et al., 2015). This raises the possibility that cortical areas may receive distinct neuromodulatory signals (Pafundo et al., 2016) and highlights a need to study neuromodulatory systems based on their projection.

We sought to elucidate the roles of neuromodulatory axons that project to the primary visual cortex. To specifically study these projections while avoiding those that solely project to other cortical areas, we used Cre recombinase driver lines and our recently described TIGRE 2.0 transgenic reporter mice to express GCaMP6s in neuromodulatory axons (Daigle *et al*., 2018). We then determined physiological and behavioral contexts that correlated with changes in the activity of these neuromodulatory projections by imaging GCaMP6-labeled axons in primary visual cortex using two-photon microscopy during periods of locomotion and pupillary movements and dilations.

### Experimental Procedures

#### Transgenic mice

To express GCaMP6s in neuromodulatory neurons, we employed three Cre driver lines and one Cre-dependent GCaMP6s reporter line:

- ChAT-Cre: B6J.ChAT-IRES-Cre::frt-neo-frt, Jax stock number 028861 (Rossi *et al*., 2011).
- Dβh-Cre_KH212, GENSAT number 032081-UCD (Gerfen *et al*., 2013).
- Slc6a4-Cre^ERT2^_EZ13 (also known as SERT-Cre^ERT2^) (Gerfen *et al*., 2013).
- Ai162(TIT2L-GCaMP6s-ICL-tTA2), Jax stock number 031562 (Daigle *et al*., 2018).

To produce mice for experiments, homozygous ChAT-Cre mice were crossed with homozygous Ai162 mice; hemizygous Dbh-Cre mice were crossed with homozygous Ai162 mice; and hemizygous Slc6a4-CreERT2 mice were crossed with homozygous Ai162 mice. We utilized the tamoxifen-inducible Slc6a4-Cre mouse line to avoid expression of Cre in thalamic and other non-serotonergic cells which transiently express SERT, and thus Cre, in early development (D’Amato *et al*., 1987). Slc6a4-Cre^ERT2^;Ai162 mice received ~200 μl of tamoxifen solution (0.2 mg/g body weight) via oral gavage once per day for five consecutive days after postnatal day 25 to activate Cre-mediated recombination. Mice were maintained on a 12-hour reverse light-dark cycle and fed *ad libitum*. All experiments were performed during the dark cycle and mice of both sexes were used for experiments. All animal procedures were approved by the Institutional Animal Care and Use Committee (IACUC) at the Allen Institute for Brain Science.

#### Histology

GCaMP6 expression was examined in two-three mice of each cross and three Ai162-only mice aged between postnatal day 60 −200. Mice were anesthetized with 5% isoflurane and intracardially perfused with 10 ml of saline (0.9% NaCl) followed by 50 ml of freshly prepared 4% paraformaldehyde (PFA) at a flow rate of 9 ml/min. Brains were rapidly dissected and post-fixed in 4% PFA at room temperature for 3-6 hours and then overnight at 4°C. Brains were subsequently incubated in 10% and then 30% sucrose in PBS for 12-24 hrs at 4°C before being cut into 50 μm sections using a freezing-sliding microtome (Leica). For histology involving the antibody-enhancement GCaMP6s-labeled neuromodulatory axons in visual cortex, we performed an antigen retrieval step consisting of heating sections to 75°C for 20 minutes in 10 mM sodium citrate pH 6.0 and 0.05% Triton-X-100.

For antibody staining, sections were collected, rinsed in PBS, and blocked with 5% normal donkey serum and 0.2% Triton X-100 in PBS for one hour. Tissue was incubated in primary antibodies diluted in the blocking solution for 48-72 hours at 4°C, washed the following day in 0.2% Triton X-100 in PBS and incubated in Alexa −488, −568, or −647 conjugated secondary antibodies (1:500, ThermoFisher) and DAPI. Sections were rinsed in 0.2% Triton X-100 in PBS followed by PBS, mounted on gelatin-coated slides and cover-slipped with prolong diamond antifade mounting media (P36965, ThermoFisher). The primary antibodies used to detect transgene expression in neuromodulatory cell- and axon-types were the anti-norepinephrine transporter (NET, 1:500, Atlas Antibodies AMAb91116), anti-tyrosine hydroxylase (TH, 1:1000, Abcam ab112), anti-serotonin transporter (SERT, 1:500, EMDMillipore MAB1564), anti-tryptophan hydroxylase 2 (TPH2, 1:250, EMD Millipore ABN60), and anti-choline acetyltransferase (ChAT, 1:300, EMDMillipore AB144P). To detect GCaMP6 in cortical axons in fixed sections, we found it necessary to enhance the GCaMP6 fluorescence with an anti-GFP primary antibody (1:5000, Abcam, Ab13970) and donkey anti-chicken secondary antibody (703-545-155, Jackson ImmunoResearch). Antibody-enhanced GCaMP6-labeled axons in cortex were imaged using a confocal microscope (FV3000, Olympus) and a 60×/1.4 NA objective. Quantification of GCaMP6 colocalization with neuromodulatory cell body markers was performed by analyzing native GCaMP6 fluorescence. Native GCaMP6 was imaged using a widefield VS120 slide loader microscope (Olympus) with a 10×/0.4 NA objective or the FV3000 confocal laser scanning microscope with a 20× objective. Neurons with colocalized GCaMP6 and a neuromodulatory cell-type marker (ChAT+, TH+, or TPH2+) were quantified using the cell counter plugin (K. De Vos, University of Sheffield) within ImageJ (FIJI).

### *In Vivo* Microscopy

#### Surgery and Habituation

Chronic cranial window implantation for long-term imaging in awake mice imaging of mouse visual cortex was performed as described previously (de Vries et al., 2018; Goldey et al., 2014). Briefly, under 0.5-2% isoflurane anesthesia, a head restraint bar was attached to the skull using C & B Metabond (Parkell) and a craniotomy was opened over the left visual cortex at coordinates 2.7 mm lateral, 1.3 mm anterior to lambda. A durotomy was performed and the craniotomy was sealed with a stack of three #1 coverslips, attached to each other using optical adhesive, and attached to the skull with Metabond. Following surgery, mice recovered for at least seven days before habituation and were housed individually.

To habituate mice to handling and head restraint, mice underwent habituation consisting of one to three days of handling by an experimenter for <10 minutes a day, followed by a day of head-restraint for <10 minutes, and then one day of head restraint next to a monitor displaying a gray screen for 10-15 minutes. Starting at the time of habituation, running wheels (InnoDome, BioServ) were placed in the cages of mice for enrichment and to encourage running behavior during imaging sessions.

#### Retinotopic Mapping

Retinotopic maps were generated through the cranial window using green wavelength reflectance in awake mice, similar to procedures described for the generation of retinotopic maps using intrinsic signal imaging (Juavinett *et al*., 2017). Images were acquired with a 1:1 optical relay using two 1× PlanAPO dissecting microscope lenses (Leica, 10450028). Illumination was from green LEDs (SuperBrightLEDS.com, NFLS-X3-LC2), via a bandpass filter (469/35, Semrock) and fluorescence was detected by a CCD camera (Flash, Hamamatsu) via a 497 nm dichroic and 525/39 bandpass filter (Semrock). Visual stimulus presentation and analysis of retinotopic maps following ISI was performed using the Retinotopic Mapping Python library of (Zhuang et al., 2017), https://github.com/AllenInstitute/retinotopic mapping. Maps were generated by sweeping a flickering black-and-white checkerboard-patterned bar across a 40” Samsung 6300 LED TV, 13.5 cm from the right eye and covering approximately −10 to 130 degrees in azimuth and −50 to 60 degrees in altitude. A gap of ≥5 s was inserted between visual stimuli presentations, resulting in repetition of the stimulus at 0.048 Hz for vertically-moving stimuli and 0.043 Hz for horizontally-moving stimuli.

#### 2-photon imaging and analysis

Axons in primary visual cortex (V1) were imaged with 2-photon excitation using a Sutter MoM and 920 nm illumination from a Ti:sapphire laser (Coherent Chameleon II), focused with a 16×/0.8 NA objective (Nikon N16XLWD-PF). We used a ~200 × 200 μm field of view with 512 × 512 pixels, acquired at 30 Hz. Emitted light was collected in the epifluorescence configuration through a 735 nm dichroic mirror (FF735-DiO1, Semrock), a 750 nm shortpass filter to remove IR laser light (FESH0750, Thorlabs) and a 470-558 nm bandpass emission filter (FF01-514/44-25, Semrock Technology). Image acquisition was controlled using Vidrio ScanImage software (Pologruto *et al*., 2003). To maintain constant immersion of the objective, we used gel immersion (Genteal Gel, Alcon). Following completion of each experiment, a Z-stack was collected from the pial surface to 250-300 microns deep in cortex. Experiments were not used if off-target, cortical cell expression was observed. Mice were head fixed on top of a rotating disk to allow them to run. During imaging, the microscope cage was continuously illuminated with a cool white (6500 K) LED (MCWHL5, Thorlabs) to partially constrict the pupils.

Fluorescence movies were motion-corrected by in the transverse optical axes using a rigid body transform based on phase correlation across frames as calculated in the OpenCV Python library. To generate axon regions of interest (ROIs), movies were down sampled in time by a factor of ten and weighted ROIs were isolated using constrained non-negative matrix factorization (CNMF) implemented in the CalmAn Python library (Giovannucci et al., 2018; Pnevmatikakis et al., 2016). The resulting ROIs were Gaussian filtered and thresholded to reduce noise. ROIs with more than 10% of their area overlap with another ROI, smaller than 150 pixels^2^, or within ten pixels from the edge of a field-of-view were excluded. For each axon ROI, a neuropil ROI was created by dilating the outer border of the axon ROI by 1 and 8 pixels to define inner and outer limits of the neuropil ROI. The union of axon ROIs was excluded from all neuropil ROIs. For each ROI we subtracted the mean fluorescence in the neuropil ROI from the mean in the axon ROI. To minimize the number of ROIs with characteristics of inactive auto-fluorescent ‘blebs’ (Reimer *et al*., 2016), we eliminated ROIs that were highly circular (eccentricity > 0.75) or which had a low skewness (<0.4) in their extracted fluorescence trace, characteristic of ROIs lacking GCaMP6 expression (Dipoppa *et al*., 2018). Time-alignment and other analyzes were performed using custom-written Python scripts and the AxonImaging library, https://github.com/Rylan-L/AxonImaging.

#### Analysis of locomotion and pupil diameter and position

To identify running epochs, we identified positive going changes in the running signal that crossed a 2 cm/s threshold and which lasted at least 1 second. The end of a running epoch was identified as a period where the running signal went below 2 cm/s for at least three seconds. In some instances, the mouse moved in the reverse direction, typically for very brief periods, and these changes were not included in our running epochs.

We monitored dilations and eye movements in both eyes by illuminating the peri-ocular region with 850 nm LED (M850LP1, Thorlabs). The image of the right eye was reflected by a low-pass dichroic mirror (Semrock, FF750-SDi02-25×36) placed ~3 cm from the eye. Images of the eyes were acquired using two cameras (Mako G-125B, Allied Vision) mounted with a zoom lens (Zoom 7000, Navitar) and an 850 nm bandpass filter (FF01-850/10-25, Semrock). Eye images were acquired at 50 Hz and running speed was acquired at 10 KHz and both signals were later down sampled to 30 Hz to match the two-photon acquisition rate.

We calculated the pupil position and diameter using custom software similar to that described previously (D. Williams) (Zhuang *et al*., 2017). Briefly, the corneal reflection of the infrared LED was first identified and masked to eliminate any potential contribution to the detection of the pupil. The image was then blurred and pupil edges were detected and exaggerated using the OpenCV package. Pupil motion in retinotopic coordinates (changes in gaze angle) was calculated under the assumption that the mouse eye is spherical, with a radius of 1.7 mm (Remtulla and Hallett, 1985). First, for each frame we determined the location of the centroid of the pupil and of the reflection of the IR-LED. From these values, we calculated the position of the pupil (in pixels) relative to the LED reflection in horizontal and vertical planes (Xi and Yi, where i represents frame number). The cardinal axes of the camera chip, which was mounted parallel to the optical table, were used to define the horizontal and vertical planes of pupil movements. Pupil positions in pixels were converted to changes in azimuth and altitude gaze angles by trigonometry: Δθazi = arcsin(ΔXi/r); Δθalt = arcsin(ΔYi/r), where ΔXi and ΔYi represent the deviation of X and Y position in the ith frame from the mean location during the movie, and r was set to be 1.7 mm (the average radius of mouse eye). The locomotion analog signal and eye movie frame timestamps were recorded on the same NI PCI 6353 digital IO board that records 2-photon image timing signal, thus enable the temporal alignment across 2-photon imaging, locomotion, and pupil recordings. We identified the baseline value for pupil diameter for each mouse by taking the median value across the entire filtered trace of the pupil diameter (low-pass Butterworth with 1 Hz cutoff). To find pupil dilations, we thresholded the unfiltered pupil diameter trace to find increases in the diameter that were 4% or more above the calculated baseline and which lasted for at least 0.5 seconds. We identified the end of a dilation by finding periods where the pupil diameter went below the threshold for at least 0.5 seconds. To find pupil dilations in the absence of running, all dilations that occurred within +/- 10 seconds the beginning or end of a running epoch were excluded.

## RESULTS

We sought to develop transgenic mouse lines that would express the genetically-encoded calcium sensor GCaMP6s (Chen et al., 2013) in neuromodulatory neuron types and would allow us to monitor the activity of the subpopulation of their axons that specifically project to the visual cortex. We previously observed non-existent or very low levels of expression in neuromodulatory neurons when we attempted to drive GCaMP6 expression in these cells with previous generations of transgenic reporter lines (Daigle et al., 2018). To overcome this, we crossed recently described TIGRE 2.0 Cre-dependent GCaMP6s (Ai162) reporter lines with ChAT-Cre, Dbh-Cre, and Slc6a4-Cre ^ERT2^ mice. This resulted GCaMP6s expression in cholinergic, noradrenergic, and serotonergic somata in basal forebrain, locus coeruleus, and dorsal raphe, respectively (Figure 1A).

**Figure 1.**
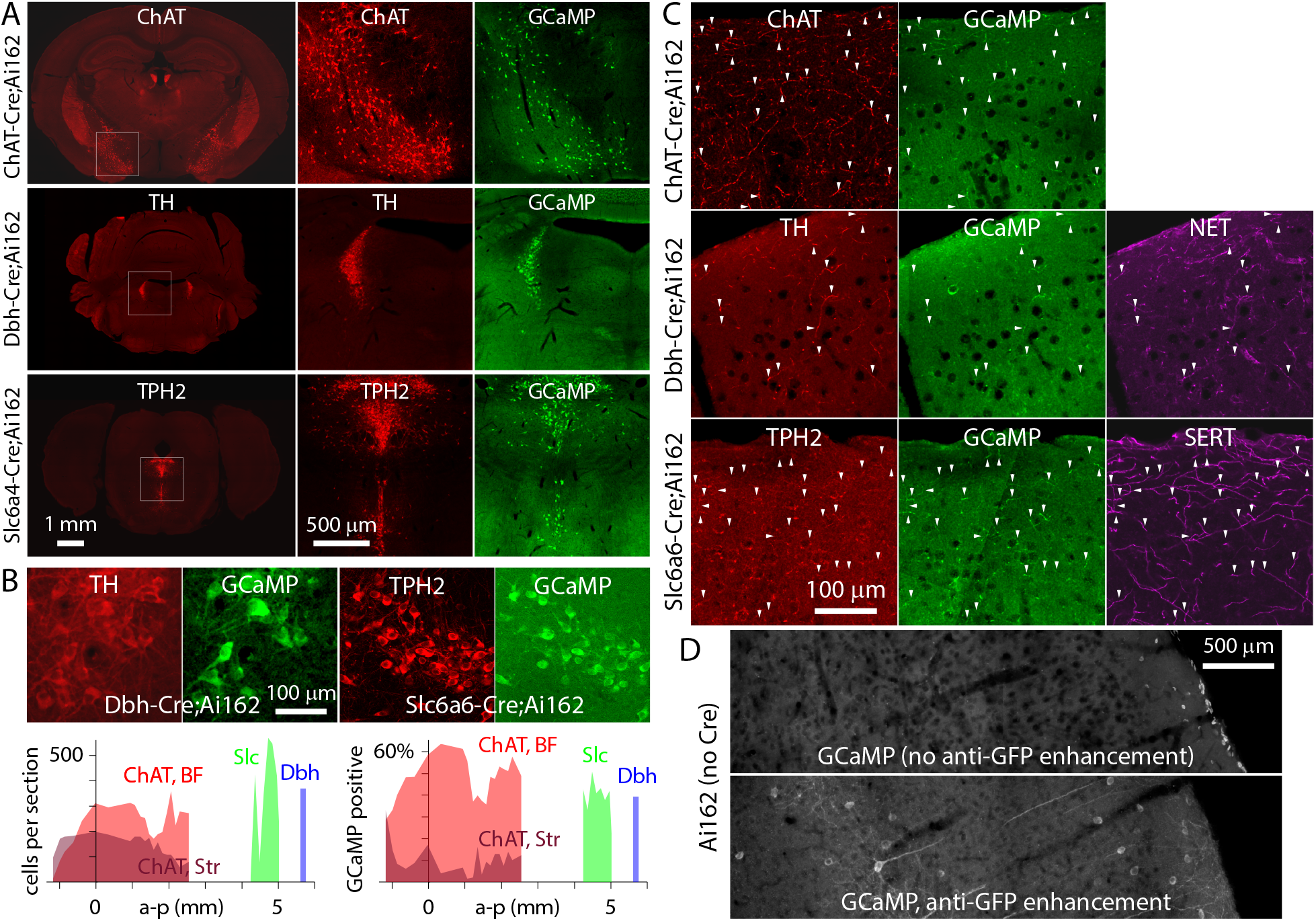
GCaMP6s expression in cholinergic, noradrenergic and serotonergic neurons and axons. (A) Widefield fluorescence images of 50 μm thick coronal sections from three mouse crosses, showing GCaMP6s expression in the basal forebrain in ChAT-Cre:Ai162 mice (n=2 mice), in locus coeruleus in Dbh-Cre;Ai162 mice (n=2 mice), and in raphe nuclei in Slc6a4-Cre^ERT2^;Ai162 mice (n=3 mice). In red, anti-choline acetyltransferase (ChAT) immunofluorescence, anti-tyrosine hydroxylase (TH) immunofluorescence, and tryptophan hydroxylase 2 (TPH2) immunofluorescence. In green, GCaMP6s fluorescence (without immuno-enhancement). (B) Confocal images of the locus coeruleus in Dbh-Cre;Ai162 mice and the dorsal raphe in Slc6a4-Cre^ERT2^;Ai162 mice, showing GCaMP fluorescence (not antibody enhanced) and anti-TH (Dbh-Cre;Ai162) or TPH2 (Slc6a4-Cre^ERT2^;Ai162). Expression of GCaMP6 was largely confined to neuromodulatory neurons in each cross with 9% of GCaMP6s-expressing somata being TH- in Dbh-Cre;Ai162 crosses (14 somata in 2 mice) and 1% of GCaMP6s-expressing somata being TPH2- (23 somata in 2 mice) Slc6a4-Cre^ERT2^;Ai162 mice. Below, soma counts per coronal section and percentage neuromodulatory somata expressing GCaMP. For ChAT-Cre;Ai162, ChAT+ neurons in the basal forebrain complex (BF) and striatum (Str) were counted separately. Expression of GCaMP6s was specific to ChAT+ neurons with 82 of 5316 GCaMP6s-expressing somata in the basal forebrain being ChAT- and 4% of GCaMP6s-expressing somata in striatum being ChAT- (19 somata). Anterior-posterior (a-p) coordinates relative to bregma. Sections were sampled at 200 μm intervals in the coronal plane. (C) Confocal images of neuromodulatory axons in the visual cortex. GCaMP-expressing axons (enhanced with anti-GFP staining, arrowheads) co-labeled with anti-ChAT in Chat-Cre;Ai162 mice, anti-TH and anti-NET (norepinephrine transporter) in Dbh-Cre;Ai162 mice, and anti-TPH2 and anti-SERT (serotonin transporter) in Slc6a4-Cre^ERT2^;Ai162 mice. (D) Native GCaMP fluorescence (no immunostaining) and anti-GFP immunostained visual cortex in coronal sections from a Ai162 reporter mouse.

In each neuromodulatory nucleus, GCaMP6s expression labeled a portion of the neuromodulatory neurons, which we confirmed by counter-staining with antibodies that were specific to each neuromodulatory neuron type (Figure 1B). In ChAT-Cre;Ai162 mice, GCaMP6s was expressed in neurons throughout the basal forebrain, with 51% (2685 of 5316 somata) of choline acetyltransferase positive (ChAT+) neurons in the basal forebrain complex expressing GCaMP6s. Colocalization of GCaMP6s in ChAT+ neurons was low in ChAT+ striatal neurons (12%, 397 of 3242 somata) and highest in the basal forebrain areas posterior to bregma that are thought to project to posterior cortices such as the visual cortex (Kim et al., 2016; Pafundo et al., 2016). GCaMP6 expression was highly specific to ChAT+ neurons with <3% of GCaMP6s-expressing somata in the basal forebrain being ChAT-. Similarly, in the locus coeruleus of Dbh-Cre;Ai162 mice, GCaMP6s was expressed in 39% (146 of 370 somata) of tyrosine hydroxylase-positive neurons (TH+). In the dorsal raphe nuclei of Slc6a4-Cre^ERT2^;Ai162 mice, GCaMP6s was expressed in 39% (2043 of 5239 somata) of tryptophan hydroxylase 2-positive neurons (TPH2+). These results demonstrate Ai162 reporter line drives expression of GCaMP6 in a portion of the neuromodulatory cells from each Cre line cross with high specificity.

To ensure that GCaMP6s was also trafficked and expressed in the cortical-projecting axons of each neuromodulatory cell type, we counterstained axons in visual cortex with neuromodulatory markers while antibody-enhancing the GCaMP6 signal to maximize our ability to detect its fluorescence in axons. Immunostaining for the axon-localized transporters, ChAT, NET, and SERT labeled cholinergic, noradrenergic, and serotonergic axon types in the primary visual cortex of each respective neuromodulatory Cre-line, and many of these axons also expressed GCaMP6s (Figure 1C). In Ai162 mice, both with and without Cre expression (reporter only), we observed that using an anti-GFP antibody to amplify GCaMP6s fluorescence unmasked small numbers cortical neurons that expressed GCaMP6s at such low levels that their fluorescence was largely undetectable in non-antibody enhanced sections (Figure 1D). These off-target labeled neurons did not colocalize with neuromodulatory markers and we excluded mice from axon imaging studies (below) if off-target, cortical GCaMP6-labeled neurons were observed *in vivo*. In conclusion, counter-staining with neuromodulatory markers indicates that crossing ChAT-Cre, Dbh-Cre and Slc6a4^ERT2^-Cre mice with the Ai162 reporter line results in largely selective expression of GCaMP6s in cholinergic, noradrenergic and serotonergic neurons and their cortical-projecting axons.

We next sought to determine behavioral conditions in which each neuromodulatory projection to the visual cortex was active. To accomplish this, we implanted mice of each Cre line with a five mm diameter cranial window over visual areas (Figure 2A) (de Vries et al., 2018; Goldey et al., 2014). To ensure that we targeted the primary visual cortex with our imaging experiments, we generated a retinotopic field sign map (Figure 2B). We used this map to target the 2-photon microscope to the center of primary visual cortex (V1), which positioned our field of view approximately on the horizontal meridian and at ~40-50° temporal from the vertical meridian in monocular V1. Fluorescence images of GCaMP6s-expressing axons in visual cortical layer 1 and upper layer 2/3 (30-100 μm below the pial surface) were acquired using 2-photon excitation (Figure 2C). During imaging, mice were free to run on a disk and we monitored both pupils for changes in position and dilation, which may correlate with changes in behavioral state (Larsen and Waters, 2018). Time-varying measurements of activity were extracted from fluorescence movies for regions of interest, each region corresponding to part of a putative axon (Figure 2D). Axons were sparse across all lines *in vivo*, with 5-69 regions per field of view (Figure 2E). However, this allowed for extraction of axonal signals from hundreds of regions of interest: 189 regions from 5 ChAT-Cre;Ai162 mice, 71 regions from 5 Dbh-Cre;Ai162 mice, and 41 regions from 3 Slc6a6-Cre;Ai162 mice.

**Figure 2.**
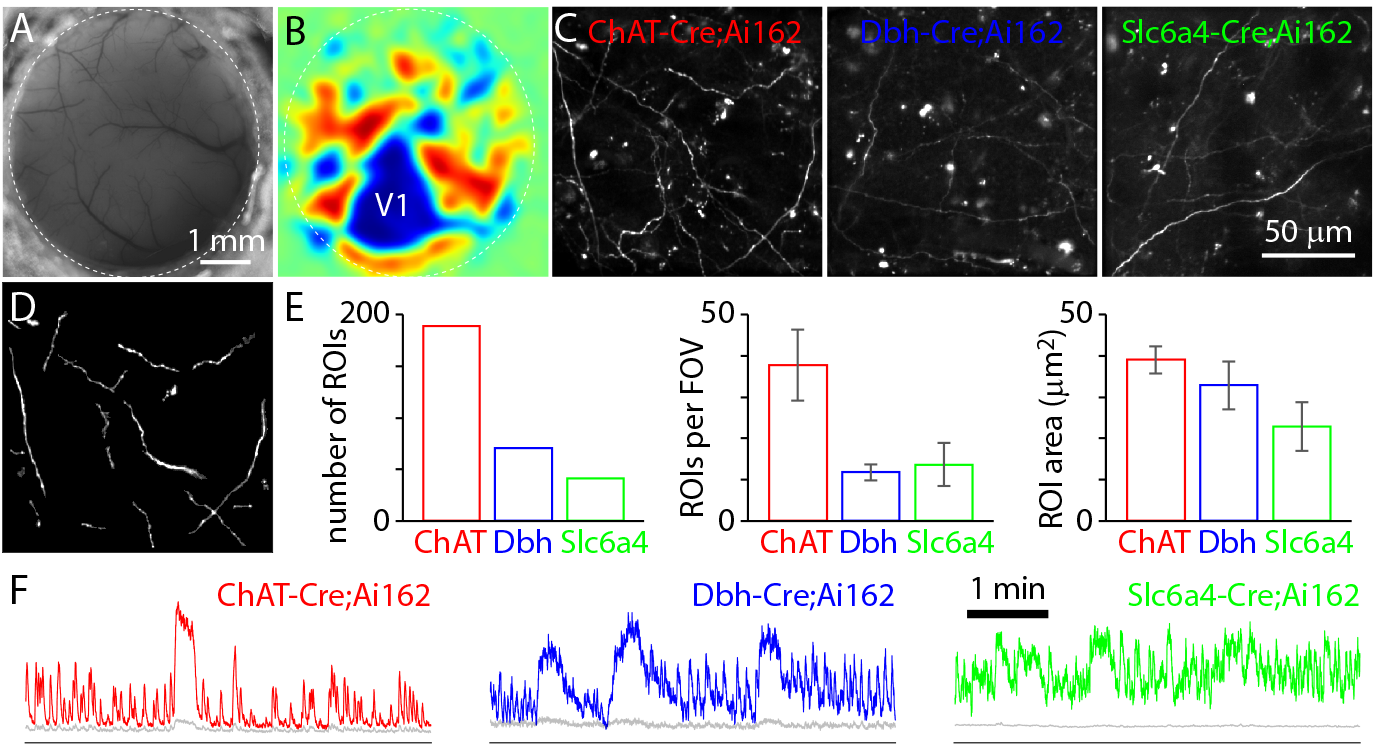
*In vivo* neuromodulatory axon imaging in primary visual cortex using GCaMP6s. (A) Example widefield image of the brain surface and five mm diameter cranial window (dashed line). (B) Retinotopic field sign map, for the window in panel A. V1: primary visual cortex. (C) Example images of axons in V1. Each is a temporal average from a single imaging plane. (D) Weighed regions of interest (ROIs) for the ChAT-Cre;Ai162 field of view in C. 24 regions of interest. (E) Mean ± SEM numbers of regions of interest and area of the regions of interest. 5 ChAT-Cre;Ai162 mice, 6 Dbh-Cre;Ai162 mice, 3 Slc6a4-Cre^ERT2^;Ai162 mice. FOV: field of view. Dashed horizontal line denotes minimum ROI size. (F) Examples of fluorescence traces. Colored lines, trace from putative axonal region of interest. Grey lines, trace from surrounding neuropil. y-axis is in arbitrary fluorescence units with the horizontal line marking zero.

Visual cortical neuron responses are modulated by locomotion (Dipoppa et al., 2018; Niell and Stryker, 2010; Saleem et al., 2013), and this is thought to increase reliability of neuronal responses to visual stimuli (de Vries et al., 2018) and decrease noise correlations (Dadarlat and Stryker, 2017). These effects have been proposed to be mediated, at least in part, by ascending cholinergic projections to the visual cortex which are activated by running (Fu et al., 2014). To determine which neuromodulatory projections were active during periods of locomotion, we imaged mice that were free to run on a disk in the absence of sensory stimuli. Many axons in ChAT-Cre;Ai162 mice exhibited increases in fluorescence, with fluorescence rising >1 s before the onset of locomotion, typically reaching a peak within ~1 s after the start of locomotion and decaying to a steady-state fluorescence during prolonged locomotion (Figure 3B). ChAT-Cre;Ai162 axon fluorescence was typically sustained throughout a running epoch and then declined to baseline after cessation of locomotion (Figure 3C). A rise in fluorescence was also observed in Dbh-Cre;Ai162 mice, again preceding locomotion by >1 s. However axon fluorescence in Dbh-Cre;Ai162 mice was often transient and declined during prolonged locomotion, sometimes to baseline. During locomotion, fluorescence in Slc6a4-Cre ^ERT2^;Ai162 axons typically did not change during running epochs.

**Figure 3.**
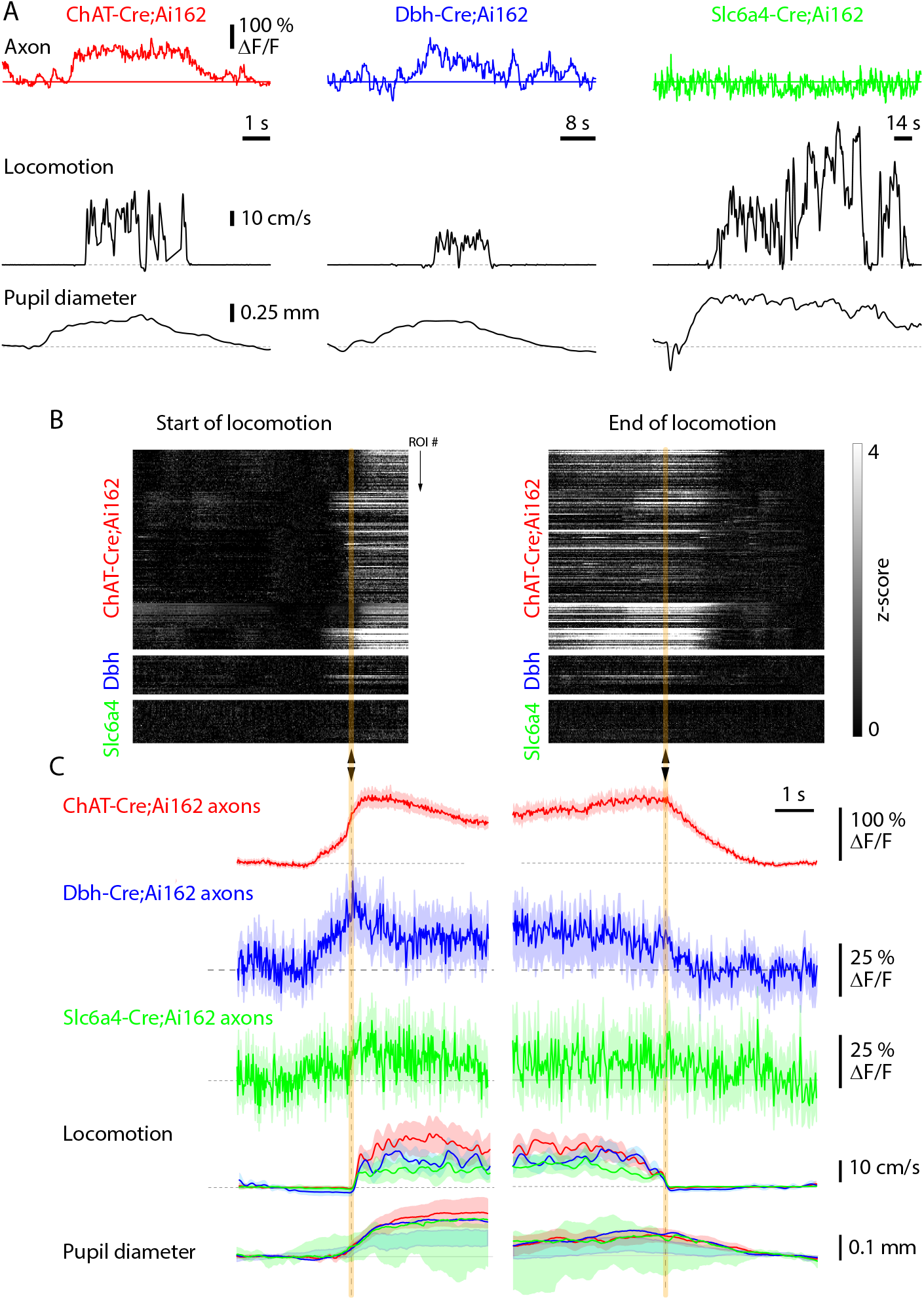
Activation of neuromodulatory axons during locomotion. (A) Activity of three example regions of interest, one from each mouse line, during a period of locomotion. Below, locomotion speed and changes in right pupil diameter. (B) Fluorescence of each region of interest at the start and end of locomotion. Each line represents the fluorescence of one region of interest during one period of locomotion. Arrow heads indicate the start and end of locomotion. n = 189, 37, and 41 axonal ROIs from ChAT-Cre;Ai162, Dbh-Cre;Ai162 and Slc6a4-Cre^ERT2^;Ai162 mice. (C) Mean (± SEM) fluorescence, locomotion and change in right pupil diameter for the three mouse lines, for ROIs with significant changes in fluorescence with locomotion. Time scale matches panel B.

Locomotion was accompanied by a large-amplitude pupil dilation that was sustained and often increased throughout the period of locomotion (Figure 4A). Between bouts of locomotion, there were also frequent, but irregularly-spaced dilations that were similar in time course to the previously described ‘microdilations’ that are associated with a transition to period of alertness (McGinley et al., 2015a). In our results, dilations in the presence and absence of locomotion (the latter we refer to as microdilations) displayed somewhat overlapping amplitude distributions, but microdilations had smaller mean peak amplitudes than locomotion-linked dilations (Figure 4B, C). Microdilations were also associated with changes in noradrenergic and cholinergic axon fluorescence (Figure 4D). The irregular occurrence of microdilations allowed us to determine whether axonal activity followed or preceded microdilations, using cross-correlation (Figure 4E). Axon activity led dilations by 0.6 s in ChAT-Cre;Ai162 mice and 0.7 s in Dbh-Cre;Ai162 mice (Figure 4G). For both axon populations, the amplitude of the fluorescence change correlated with amplitude of the dilation (Figure 4F, H).

**Figure 4.**
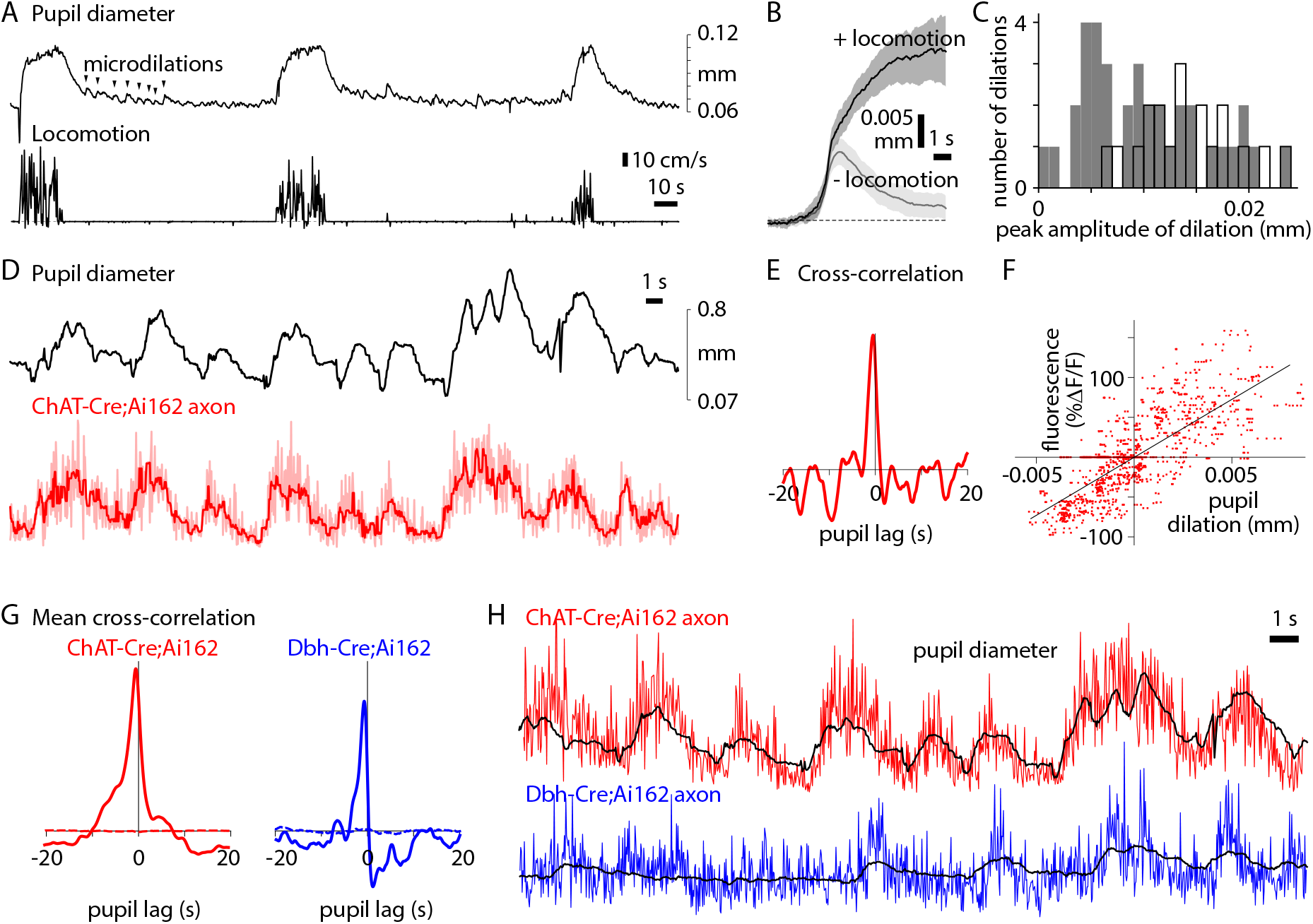
Pupil dilation-linked changes in neuromodulatory axon fluorescence. (A) Pupil dilations and locomotion over several minutes in a ChAT-Cre;Ai162 mouse. Microdilations (arrow heads, examples) often occurred in bouts between periods of locomotion. (B) Mean (± 95% confidence intervals) change in pupil diameter for pupil dilations with and without locomotion. (C) Histogram showing distribution of amplitudes for dilations with and without locomotion. (D) Microdilations occurring over 40 seconds, accompanied by changes in axon fluorescence (red) in a ChAT-Cre;Ai162 mouse. Dark red, filtered fluorescence trace. There was no locomotion during this period. (E) Cross-correlation of the pupil and filtered fluorescence traces in panel B. Peak is at < 1 s, indicating that pupil dilation lagged the fluorescence signal. (F) Relationship between fluorescence and pupil diameter for the example in panel B. Linear fit. (G) Average cross-correlations for ChAT-Cre;Ai162 and Dbh-Cre;Ai162 mice. Dashed lines, mean cross-correlation after shuffling (in the temporal axis) the fluorescence trace for each dilation event. (H) Example traces, showing that fluorescence (color) and pupil diameter (black) were closely matched after temporally advancing the fluorescence trace and scaling by the mean slope of the relationship between fluorescence and pupil diameter.

In addition to pupil dilations, mice perform saccadic-like eye movements, typically in the horizontal (azimuth) direction, during exploration (Samonds et al., 2018). Because periods of locomotion were preceded by large-amplitude pupil dilations (Figure 4A) (McGinley et al., 2015a; Reimer et al., 2014), we next sought to determine if locomotion itself increased axon fluorescence in ChAT-Cre;Ai162 and Dbh-Cre;Ai162 mice, or if running increases in fluorescence could be solely attributed to preceding pupillary dilations or movements. To address this, we compared pupil dilation-triggered average traces for dilations that occurred with and without locomotion while measuring horizontal and vertical position eye position of the left and right eyes (Figure 5A). In ChAT-Cre;Ai162 mice, the mean pupil dilation that preceded the onset of locomotion was larger in amplitude and more sustained than dilations in the absence of locomotion, suggesting locomotion additionally contributed to the magnitude of the dilation (McGinley et al., 2015a). However, in Dbh-Cre;Ai162 mice, pupil dilations were similar in the presence and absence of locomotion. Based on this, we created a predicted fluorescence trace by time shifting and scaling the mean pupil diameter. In the absence of locomotion, the pupil-predicted fluorescence matched the amplitude and time course of axon fluorescence in ChAT-Cre;Ai162 and Dbh-Cre;Ai162 mice (Figure 5B). In trials with locomotion, the pupil-predicted fluorescence trace accounted for much, but not all, of the measured fluorescence changes. In ChAT-Cre;Ai162 mice and Dbh-Cre;Ai162 mice, the predicted and measured fluorescence traces matched as locomotion speed increased and peaked, but thereafter the predicted fluorescence exceeded the measured fluorescence. Our results suggest that much of the locomotion-associated activity of cholinergic and noradrenergic axons is related to the pupil dilation that precedes locomotion. However, the mismatch in measured and pupil-predicted changes in fluorescence suggests that axon activity is suppressed relative to the continued dilation of the pupil after the initial phase of locomotion. Additionally, locomotion was associated with dilations of both pupils and eye movements, with both pupils moving nasally (forward) as the pupils dilated (Figure 5A). In the absence of locomotion, right pupil dilation was typically accompanied by dilation of the left pupil and sometimes by small nasal (forward) movements of both pupils (Figure 5A), but these eye movements were more pronounced during locomotion. In summary, our results demonstrate that the axon fluorescence in ChAT-Cre;Ai162 and Dbh-Cre;Ai162 increases during periods of pupillary dilation and movements, both when mice are stationary or running.

**Figure 5.**
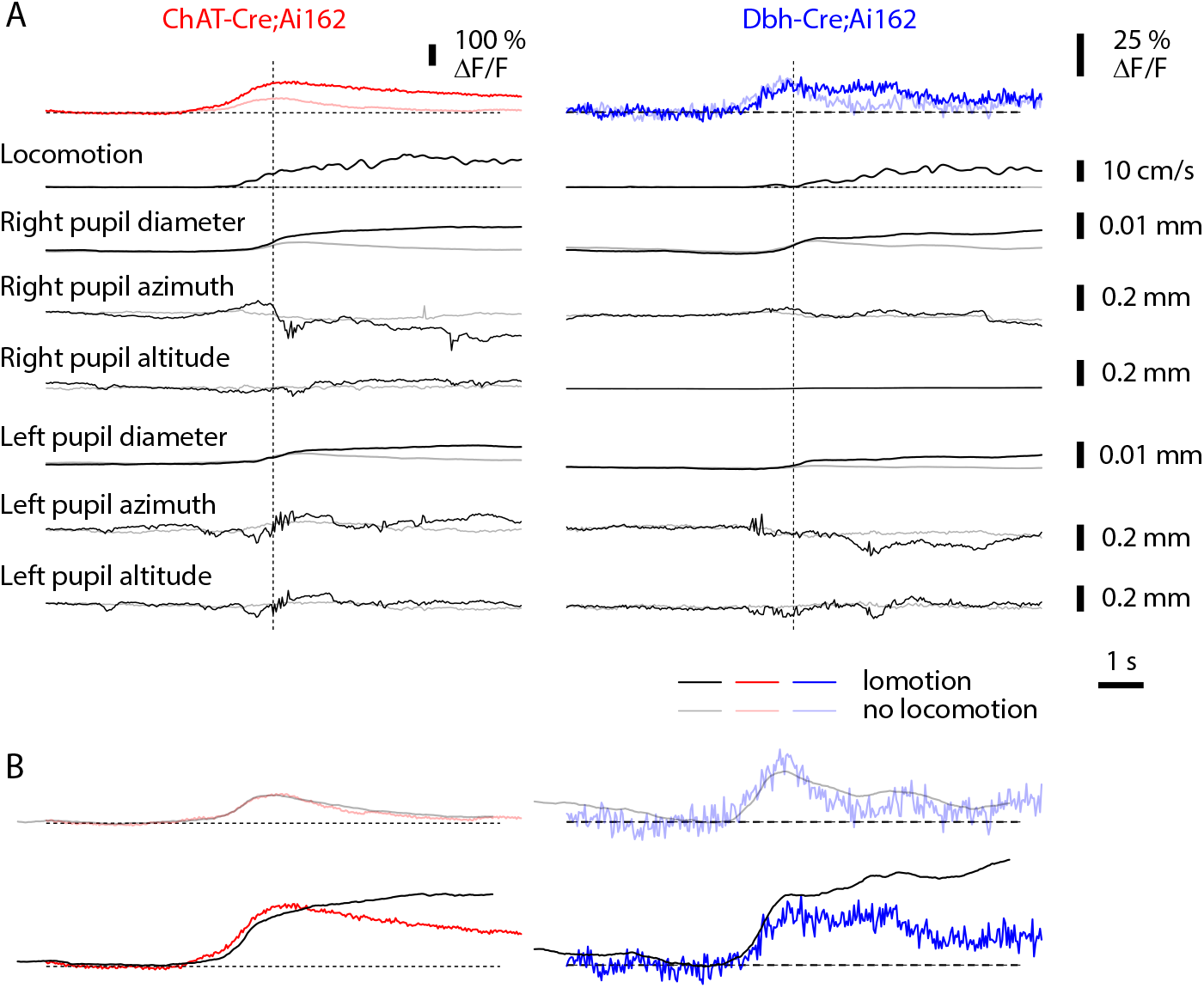
Relationship between pupil-linked axon fluorescence and locomotion. (A) Fluorescence, pupil diameter, and pupil movements in horizontal (azimuth) and vertical (altitude) directions during stationary periods or during locomotion. Each line is an average across all ROIs and trials in the top fluorescence trace, with pupil dilation epochs triggered off changes in the right pupil diameter. (B) Predicted fluorescence traces based on the magnitude of the pupil dilation in ChAT-Cre;Ai162 (red) and Dbh-Cre;Ai162 mice (blue).

## DISCUSSION

We have described several Cre-driver x TIGRE 2.0 GCaMP6 reporter mouse line crosses that drive expression of GCaMP6s in neuromodulatory neurons and their cortical-projecting axons. Previous transgenic lines either failed to express GCaMP calcium sensors in neuromodulatory populations or expressed them at levels too low to support axon imaging (Daigle *et al*., 2018). This perhaps results from high somatic concentrations being required to traffic sufficient GCaMP6 sensor along these long axons, which were often millimeters from the somata in our experiments (Li *et al*., 2018).

Our results demonstrate that cholinergic and noradrenergic axon projections to the visual cortex are active before locomotion and pupil dilation. The fluorescence changes in these two axon populations displayed different time courses, with noradrenergic axons being transiently active just before the onset of pupil dilation or locomotion and cholinergic axons exhibiting more sustained activity during these periods. Our results are consistent with previous results demonstrating that cholinergic axon projections to the cortex are activated during transitions to active, exploratory behavioral states including whisking (Eggermann et al., 2014; Reimer et al., 2014), locomotion (Fu et al., 2014; McGinley et al., 2015a), and pupil dilation (Reimer *et al*., 2016).

In our experiments, locomotion was invariably accompanied by pupil dilation, in agreement with previous results (McGinley et al., 2015a; Reimer et al., 2014). We also found that locomotion is accompanied by nasally-oriented eye movements, which were similar to previously described mouse saccades (Samonds et al., 2018). Thus, pupil dilation and pupil movement are both correlated with neuromodulatory axon activity in cortex prior to locomotion. Our results suggest that much of the locomotion-linked axon activity is from pupil dilation, but that pupil dilation is not the only contributory factor. Because the magnitude axonal fluorescence seems to be modulated by pupillary changes and then additionally by running, this suggests cholinergic neurons may integrate behavioral state-associated signals from these two systems.

We demonstrate that the Slc6a4-Cre^ERT2^ x Ai162 (TITL2-GCaMP6s) mouse cross allows for the imaging of serotonergic axons in visual cortex. A subset of spinal cord-projecting serotonergic axons are thought to control locomotion initiation (Cabaj *et al*., 2017), but the activity of serotonergic axons in cortex during locomotion, pupil dilation and eye movements had previously not been explored. Serotonergic axons displayed spontaneous changes in fluorescence, suggesting that GCaMP6s was functional in serotonergic axons, but we observed little or no locomotion- or pupil-driven responses of these axons. Further experiments will be required to determine whether the lack of locomotion- and pupil-linked signals reflect a lack of activity in serotonergic axons or insufficient signal-to-noise ratio in our measurements.

At the onset of locomotion, the membrane potentials of cortical pyramidal neurons depolarize at around the same time as cholinergic and noradrenergic axon activity, ~one second before pupil dilation (McGinley et al., 2015a), raising the possibility that release of neuromodulators leads to activation of pyramidal neurons. There are likely to be other effects of neuromodulators in cortex and these effects will probably depend on postsynaptic cell type (Fu et al., 2014; Kalmbach et al., 2012), but the overall effect of neuromodulation in sensory cortices may be to change sensory processing. For example, visual cortical neurons become more responsive to their preferred orientations of drifting grating during pupil dilation (Reimer *et al*., 2014) and periods of locomotion are associated with a decrease in noise correlations between neurons, which is thought to enhance stimulus discriminability (Dadarlat and Stryker, 2017). Our results provide evidence that the axons of multiple neuromodulatory systems are active before and during some of these behavioral state changes. However, additional mechanistic studies are still needed to further understand the causal relationships between behavioral state, neuromodulation, and cortical function.

## Acknowledgements

We wish to thank Christof Koch for helpful suggestions and edits to our manuscript. Our work was funded by the Allen Institute for Brain Science and supported by award number R01NS078067 from the National Institute of Mental Health. This manuscript’s contents are solely the responsibility of the authors and do not necessarily represent the official views of the National Institutes of Health and the National Institute of Neurological Disorders and Stroke. GCaMP constructs were provided by the Howard Hughes Medical Institute, Janelia Research Campus. We thank the many staff members of the Allen Institute especially the In Vivo Sciences team for surgery assistance, transgenic colony management, genotyping, and tissue preparation. We also wish to thank Derric Williams for developing eye tracking and acquisition software. We wish to thank the Allen Institute founder, Paul G. Allen for his vision, encouragement and support.

